# Mutation of a highly conserved isoleucine residue in loop 2 of several *β*-coronavirus macrodomains indicates that enhanced ADP-ribose binding is detrimental to infection

**DOI:** 10.1101/2024.01.03.574082

**Authors:** Catherine M. Kerr, Jessica J. Pfannenstiel, Yousef M. Alhammad, Joseph J. O’Connor, Roshan Ghimire, Rakshya Shrestha, Reem Khattabi, Pradtahna Saenjamsai, Srivatsan Parthasarathy, Peter R. McDonald, Philip Gao, David K. Johnson, Sunil More, Anuradha Roy, Rudragouda Channappanavar, Anthony R. Fehr

**Affiliations:** Department of Molecular Biosciences, University of Kansas, Lawrence, Kansas 66045, USA; Department of Veterinary Pathobiology, Oklahoma State University, Stillwater, Oklahoma 74078, USA; High Throughput Screening Laboratory, University of Kansas, Lawrence, Kansas 66047, USA; Protein Production Group, University of Kansas, Lawrence, Kansas 66047, USA; Molecular Graphics and Modeling Laboratory and the Computational Chemical Biology Core, University of Kansas, Lawrence, Kansas 66047, USA; Oklahoma Center for Respiratory and Infectious Diseases, College of Veterinary Medicine, Stillwater, Oklahoma 74078, USA

**Keywords:** coronavirus, MERS-CoV, SARS-CoV-2, MHV, non-structural protein 3, macrodomain, ADP-ribosylation, ADP-ribosylhydrolase, interferon

## Abstract

All coronaviruses (CoVs) encode for a conserved macrodomain (Mac1) located in nonstructural protein 3 (nsp3). Mac1 is an ADP-ribosylhydrolase that binds and hydrolyzes mono-ADP-ribose from target proteins. Previous work has shown that Mac1 is important for virus replication and pathogenesis. Within Mac1, there are several regions that are highly conserved across CoVs, including the GIF (glycine-isoleucine-phenylalanine) motif. To determine how the biochemical activities of these residues impact CoV replication, the isoleucine and the phenylalanine residues were mutated to alanine (I-A/F-A) in both recombinant Mac1 proteins and recombinant CoVs, including murine hepatitis virus (MHV), Middle East respiratory syndrome coronavirus (MERS-CoV), and severe acute respiratory syndrome coronavirus 2 (SARS-CoV-2). The F-A mutant proteins had ADP-ribose binding and/or hydrolysis defects that led to attenuated replication and pathogenesis in cell culture and mice. In contrast, the I-A mutations had normal enzyme activity and enhanced ADP-ribose binding. Despite increased ADP-ribose binding, I-A mutant MERS-CoV and SARS-CoV-2 were highly attenuated in both cell culture and mice, indicating that this isoleucine residue acts as a gate that controls ADP-ribose binding for efficient virus replication. These results highlight the function of this highly conserved residue and provide unique insight into how macrodomains control ADP-ribose binding and hydrolysis to promote viral replication.

**AUTHOR SUMMARY:** The conserved CoV macrodomain (Mac1) counters the activity of host ADP-ribosyltransferases by removing ADP-ribose from target proteins and is critical for CoV replication and pathogenesis. Mac1 is a potential therapeutic target for CoV disease and several groups are actively developing Mac1 inhibitors. However, we lack a basic knowledge of how many of the key residues in the Mac1 ADP-ribose binding pocket contribute to its biochemical and virological functions. In this study, we engineered alanine mutations into two highly conserved residues in the ADP-ribose binding pocket of Mac1, both as recombinant proteins and recombinant viruses for both MERS-CoV and SARS-CoV-2 to determine their importance in both Mac1 biochemical functions and CoV infection. Interestingly, an isoleucine-to-alanine mutation in loop 2 of both MERS-CoV and SARS-CoV-2 Mac1 proteins enhanced ADP-ribose binding. But surprisingly, that proved to be detrimental to virus infection, indicating that this isoleucine functions to control Mac1 ADP-ribose binding and is beneficial for virus replication and pathogenesis. These results provide unique insight into how macrodomains control ADP-ribose binding to promote infection and will be critical for the development of novel inhibitors targeting Mac1 that could be used to treat CoV-induced disease.

## INTRODUCTION

Coronaviruses (CoVs), from the family *Coronaviridae* of the order *Nidovirales*, are large positive-sense RNA viruses of both human and veterinary significance. In the past few decades, there have been three significant outbreaks of lethal human CoV disease. The first outbreak occurred in 2002-2003 when severe acute respiratory syndrome (SARS-CoV) emerged in China. In 2012, Middle East respiratory syndrome (MERS)-CoV was reported in Saudi Arabia. More recently, in late 2019, the SARS-CoV-2 emerged in Wuhan, China and rapidly spread around the world, becoming the first CoV to be the cause of a pandemic (1).

CoVs genomes range in size from 26-32 kb and encode for four conserved structural proteins, up to ten accessory proteins, and 15-16 non-structural proteins (nsps) that are expressed in two polyproteins, polyprotein 1 a (pp1a) and polyprotein 1 ab (pp1ab). These polyproteins are further cleaved by viral proteases into individual non-structural proteins. The nsps encode for a variety of proteins that are necessary for replication and innate immune evasion, such as the RNA-dependent RNA polymerase, main protease, helicase, N-7 methyltransferase, exoribonuclease, an endoribonuclease, and many others. The largest of the CoV nsps is nsp3, which encodes for multiple modular domains, including ubiquitin-like domains, nucleic acid binding domains, one or two papain-like protease domains, a deubiquitinase domain, a CoV-Y domain, and one or more macrodomains (2, 3). Nsp3 is also required for double-membrane vesicle (DMV) formation and forms a pore within DMVs in the replication-transcription complex (RTC). Several domains of nsp3 are required to form the pore within the RTC, including those from Ubl2 to the C-terminus of nsp3 (4, 5).

Some CoVs encode for as many as 3 tandem macrodomains, termed Mac1, Mac2, and Mac3. Mac1 is a conserved domain found in all CoVs, Togaviruses, Rubiviruses, and Hepatitis E virus (6). The highly conserved structure of Mac1 consists of several central β-sheets surrounded by 3 α-helices on each side (7–11). Mac1 has been shown to reverse ADP-ribosylation of proteins *in vitro* (12–17). ADP-ribosylation is a common posttranslational modification that is catalyzed by ADP-ribosyltransferases (ARTs) that utilize NAD^+^ to covalently attach a single ADP-ribose unit, mono-ADP-ribosylation (MARylaytion/MAR), or a chain of several ADP-ribose units, poly-ADP-ribosylation (PARylation/PAR) to a target protein or nucleic acid (18, 19). These modifications are crucial for cellular stress responses, including viral infections (20). The most common mammalian ARTs are PARPs, and the most well studied PARP is PARP1, which is a target of anti-cancer drugs due to its role in the DNA damage response. However, most PARPs are mono-ADP-ribosyltransferases, many of which are IFN-stimulated genes (ISGs) and are known to impact virus infections (21).

Several studies have demonstrated that the conserved macrodomain is critical for CoV, alphavirus, and Hepatitis E virus infection, using either deletion or point mutations of the conserved macrodomain (12, 22–30), indicating that these viruses are especially sensitive to host PARP-mediated antiviral functions. However, despite clear evidence that Mac1 is critical for CoV infection, there have only been a few studies that have investigated how individual residues contribute to the biochemical functions of Mac1 and how those biochemical activities correlate with CoV replication and pathogenesis. Initial studies of Mac1 in CoVs focused on the mutation of a highly conserved asparagine residue to alanine or aspartic acid. This mutation largely impairs Mac1 deMARylating activity, as has been demonstrated for the *⍺*-CoV 229E, and the *β*-CoVs SARS-CoV, and SARS-CoV-2 proteins (9, 12, 29, 31). In most cases, these mutations did not significantly reduce virus replication in cell culture but did lead to highly attenuated viruses in mice (12, 25, 26, 29, 31). We also showed that the Asn-to-Ala mutation in the *β*-CoV murine hepatitis virus strain JHM (MHV) led to poor replication in primary bone-marrow-derived macrophages, which was reversed upon either knockdown or knockout of PARP12, demonstrating that Mac1 specifically counters PARP activity (32, 33). Additionally, the Asn-to-Ala mutation in MHV and SARS-CoV, as well as a complete Mac1 deletion and an Asn-to-Asp mutation in SARS-CoV-2, lead to increases in IFN production following infection (12, 24, 29, 32). This function was independent of its effect on virus replication, as the MHV Asn-to-Ala virus remained attenuated in MAVS^-/-^ cells and mice (32). This increase in IFN production following an MHV Mac1 mutation infection, specifically the N1347A infection, is due to its ability to counter PARP activity, as PARP inhibitors abolish this response (32). In addition, virus with the same mutation in SARS-CoV-2 (N1062A), as well as a complete Mac1 deletion virus, was more sensitive to IFNγ treatment in cell culture and was attenuated *in vivo* when compared to WT virus (24). Furthermore, overexpression of SARS-CoV-2 WT Mac1, but not the Asn-to-Ala mutation, reverses PARP14/PARP9/DTX3L mediated ADP-ribosylation following IFNγ or poly(I:C) treatment (Hoch/Ahel papers) (16). The only other mutations that have been evaluated both in the context of biochemical activity and virus replication are D1022A, H1045A, and G1130V (all numbers based on the SARS-CoV Mac1) (12, 31). In the context of SARS-CoV, each of these mutations had reduced ADP-ribosylhydrolase activity, replicated normally in cell culture, and were attenuated in mice (12). A later study evaluated the role of the D1329A and G1439V mutations in MHV. Recombinant viruses with those mutations, which are predicted to dramatically reduce Mac1 ADP-ribose binding, were highly attenuated in cell culture, much more so than the aforementioned N1347A mutation (34). These results indicate that ADP-ribose binding may be more critical for MHV than for SARS-CoV. Additional evidence supporting this hypothesis was obtained from SARS-CoV-2, as the orthologous D1044A mutation did not display increased sensitivity to IFNγ and was only partially attenuated in mice (24).

The highly conserved isoleucine and phenylalanine residues in loop 2 are also of great interest due to their conservation and positioning within the ADP-ribose binding pocket (Fig. 1A-B) (7, 9, 11), though the impact of these residues on CoV replication has not been addressed. The isoleucine appears to interact with the final glycine in loop 1 to form a narrow channel around the diphosphate within the ADP-ribose binding domain that could be important for both binding and hydrolysis. In contrast, the phenylalanine residue makes van der Waals interactions with the distal ribose and orients the distal ribose for hydrolysis. In prior studies, mutation of phenylalanine-to-leucine reduced enzyme activity of SARS-CoV-2 Mac1 as expected, while an isoleucine-to-alanine mutation of the SARS-CoV-2 or HKU4 Mac1 did not impact enzyme activity (9, 35). It remains unclear how these residues impact ADP-ribose binding and ultimately how their biochemical functions relate to virus replication and pathogenesis.

**Figure 1.**
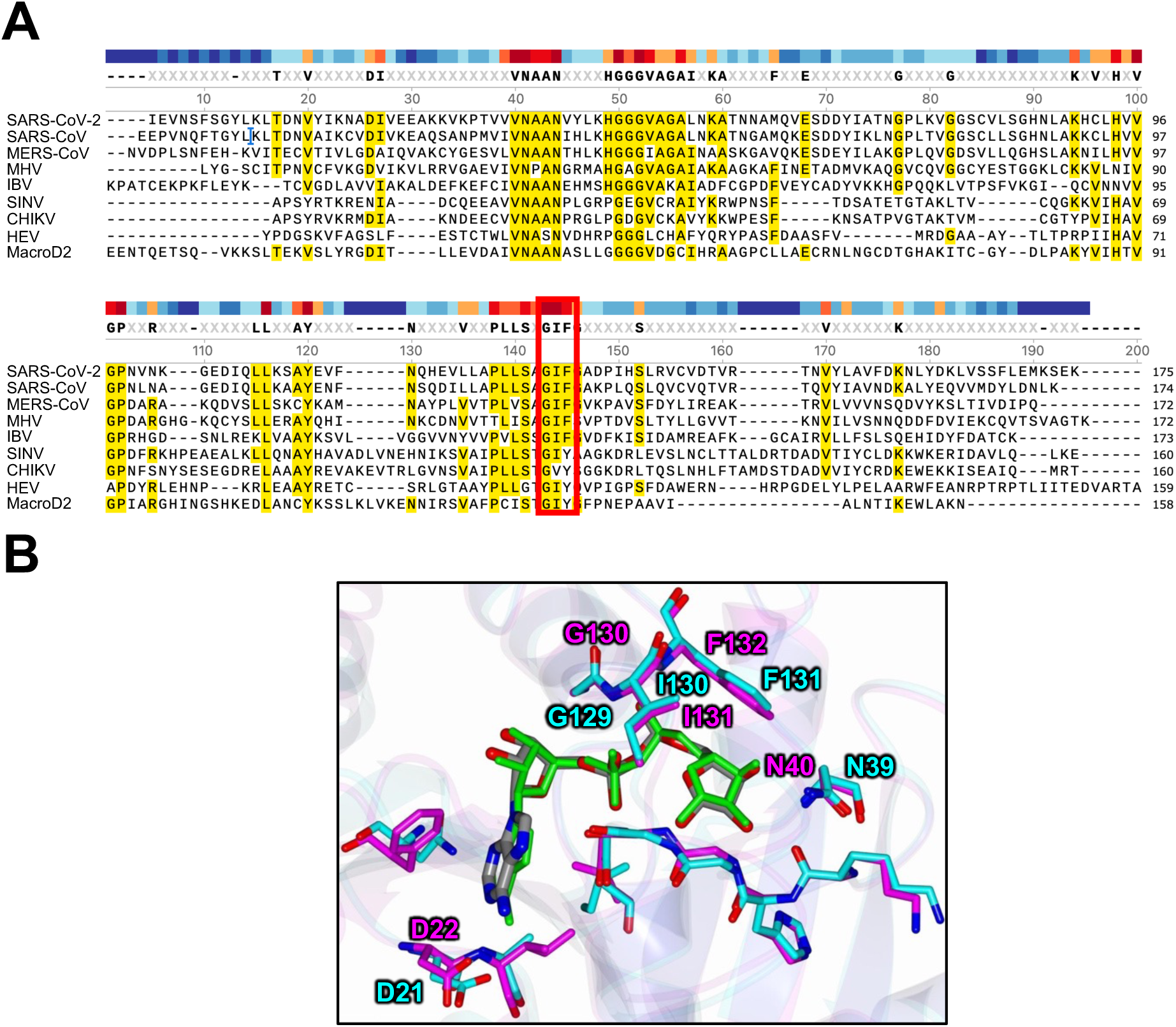
The GIF motif in loop 2 of Mac1 is highly conserved and is closely associated with both phosphate groups and the terminal ribose of ADP-ribose. (A) Sequence alignment of Mac1 across viral and human macrodomains. The GIF motif is boxed in Red. (B) Overlay of the SARS-CoV-2 (purple) (6WOJ) and MERS-CoV (teal) (5HOL) Mac1 ADP-ribose binding domains with ADP-ribose (7), highlighting the GIF motif and conserved asparagine and aspartic acid residues discussed in the manuscript.

In this study, we tested how mutations of the isoleucine (I) and phenylalanine (F) residues in loop 2 of the CoV Mac1 (Fig. 1A-B) to alanine (A) impact both Mac1 biochemical functions and the replication of *Embeco*, *Merbeco*, and *Sarbeco* lineages of *β*-CoVs. While F-A mutations resulted in poor enzyme activity and attenuated viruses, I-A mutations surprisingly led to enhanced ADP-ribose binding with little-to-no effect on its enzyme activity. More remarkably, this enhanced binding was detrimental to its biological function, as recombinant MERS-CoV and SARS-CoV-2 viruses with the I-A mutation were highly attenuated. We hypothesize that this isoleucine residue acts as a gate to control ADP-ribose binding and maintain proper enzyme activity during infection. These results provide a unique example where enhancing the biochemical activity of a protein has a negative impact on its biological function.

## RESULTS

### Mouse hepatitis virus strain JHM (MHV) F1441A, but not I1440A, has decreased replication in cell culture and in mice

To uncover the relative contributions of the residues in the highly conserved GIF motif of Mac1 in CoV replication and pathogenesis, we first compared the replication of the *Embecovirus* MHV I1440A and F1441A mutations to WT and a previously characterized mutant virus, N1347A. Previously, we found that N1347A replicates normally in most cell lines susceptible to MHV but replicates poorly in primary macrophages and in mice (26, 32). Here, we tested the replication of recombinant viruses in several different cell types that are susceptible to MHV including: a mouse astrocyte cell line (DBT), a mouse fibroblast cell line (L929), and primary bone-marrow cells differentiated into M2 macrophages. M2 macrophages were used as they have a more pronounced phenotype with the MHV N1347A virus than M0 macrophages, making them an ideal cell type to study mutant virus replication. As expected, the F1441A mutation had decreased production of infectious virus in all cell types and in mice, with defects seen at peak titers of 2.7-fold in DBT cells, 4.3**-**fold in L929 cells, and 21.7-fold in M2 macrophages (Fig. 2A-C). We previously found that a knockout (KO) of PARP12 can restore the infectious virus production of N1347A, therefore we also tested the ability of F1441A to replicate in PARP12 KO M2 BMDMs (Fig. 2D). In the absence of PARP12, F1441A infectious virus production increased by 9.3-fold, indicating that PARP12 contributes to the restriction of this virus, much like N1347A. In contrast, I1440A produced virus at WT levels in all cell types and at all time points (Fig. 2A-D).

**Figure 2.**
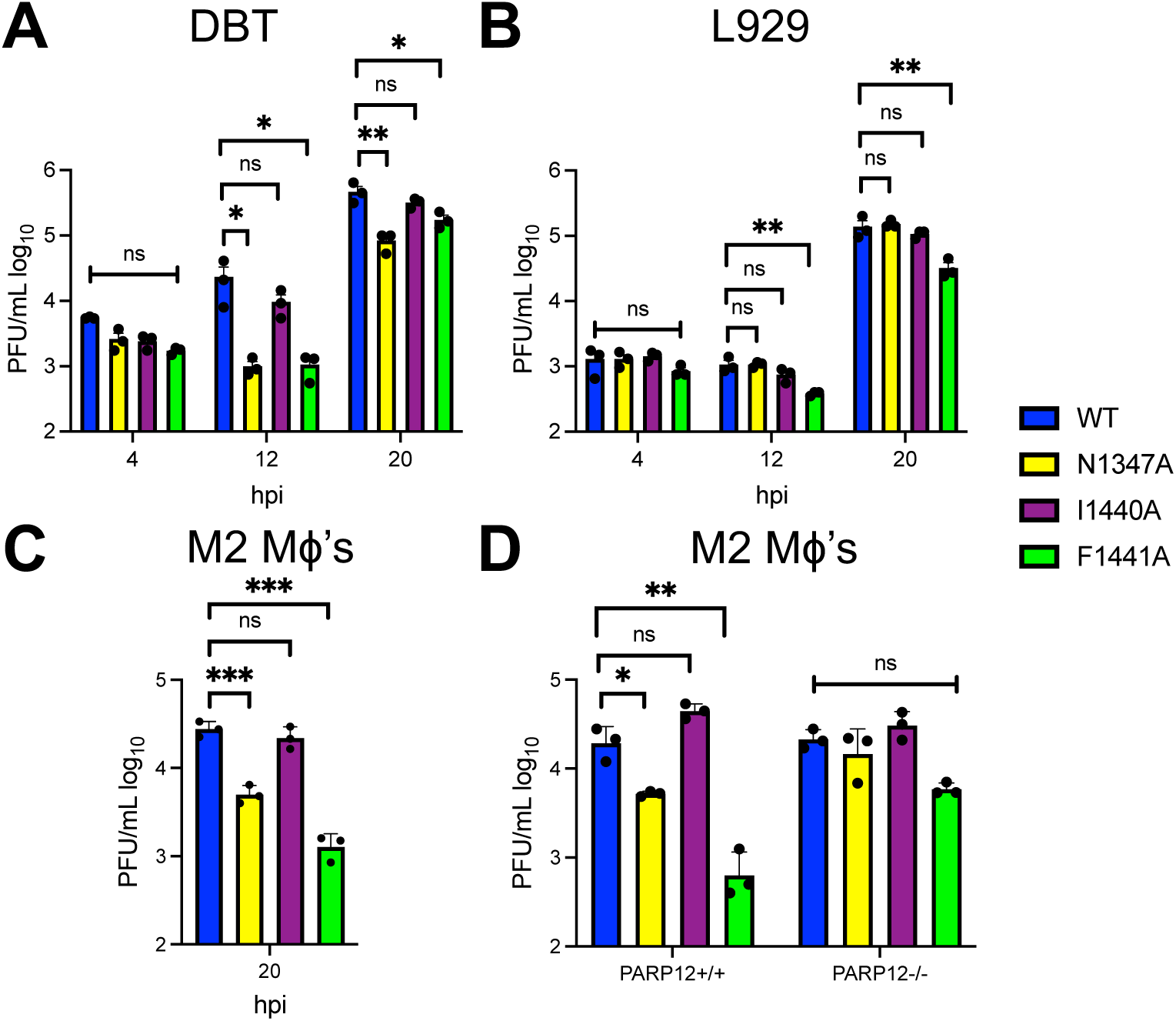
MHV F1441A mutation is attenuated in cell lines and in primary cells. DBTs (A), L929s (B), and M2 macrophages (C) were infected with JHMV at an MOI of 0.1 PFU/cell. Cells and supernatants were collected at indicated times and assayed for progeny infectious virus by plaque assay. The data in each panel show one experiment representative of three independent experiments with n = 3 for each experiment.

We hypothesized that the importance of the isoleucine residue may become more apparent *in vivo*, so we tested the ability of these MHV mutant viruses to replicate and cause severe encephalitis in mice. C57BL/6 mice were infected intranasally with 10^4^ PFU of each virus and were monitored for weight loss and survival over 12 days, and viral loads in the brain were measured at day 5 post-infection. The F1441A mutant was attenuated in mice as only 50% of the mice succumbed to infection, while the other half recovered after losing ∼10% of their body weight (Fig. 3A and B). This attenuation of F1441A was also demonstrated in the disease scores, as they began to return to normal by day 8 (Fig. 3C). Furthermore, F1441A virus infected mice had ∼7.5 fold lower viral loads than WT virus infected mice (Fig. 3D). These titers were highly variable, reflecting the fact that 50% of the mice survived. In contrast, I1440A infected mice all succumbed to disease by 9 dpi, and much like the cell culture results the I1440A viral loads were nearly equivalent to WT virus in mice (Fig. 3A-D). Taken together, this indicates that F1441 is required for efficient virus replication and disease progression, while mutation of the I1440 residue does not impact MHV replication or pathogenesis. The lack of any impact of the I1440A mutation on MHV replication or pathogenesis was surprising, considering the extreme conservation of this residue through all CoVs (6).

**Figure 3.**
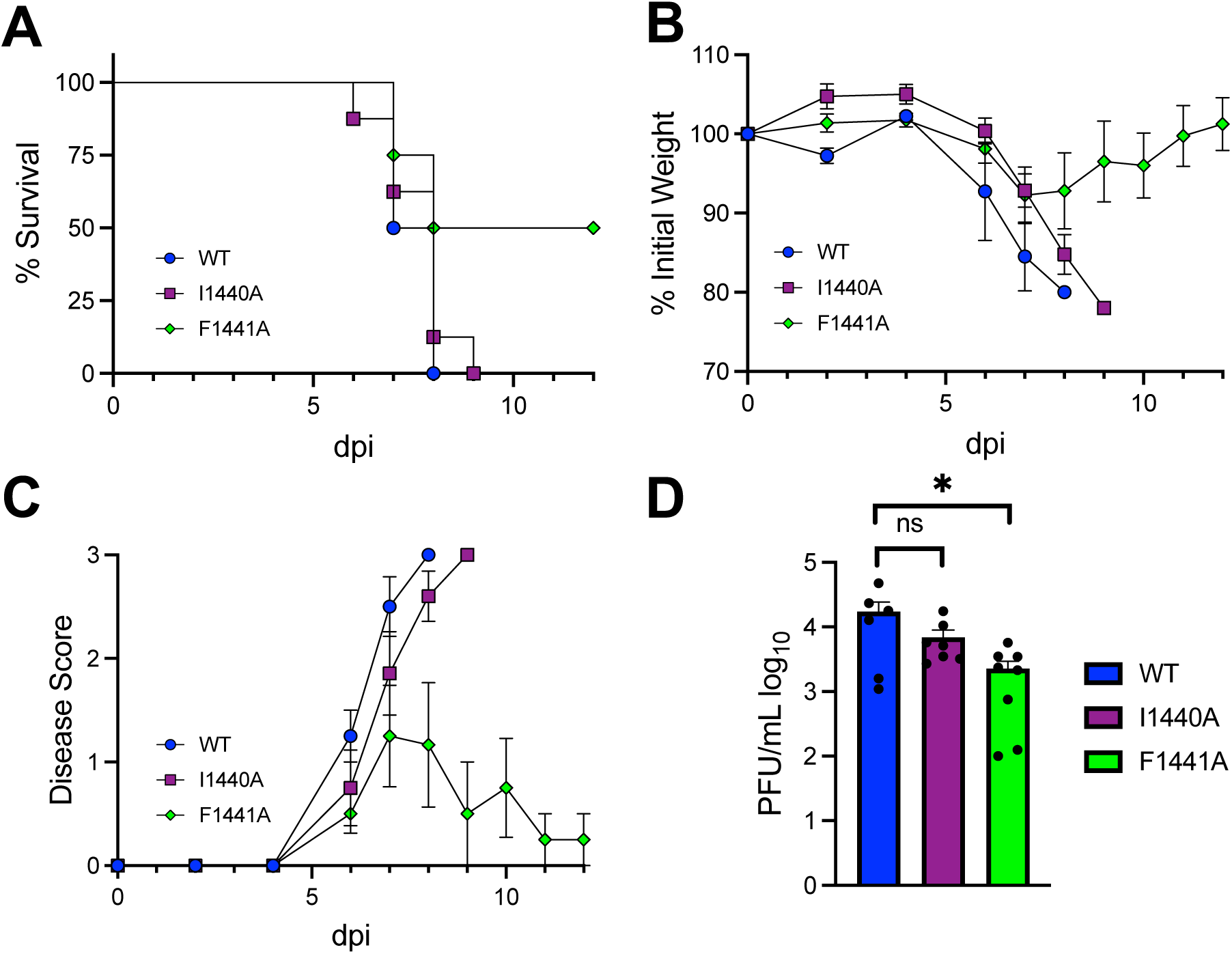
MHV F1441A, but not I1440A, is partially attenuated in *in vivo*. (A-C) Male and female C57BL/6 mice were infected intranasally with WT, I1440A, and F1441A JHMV at 1 × 10^4^ PFU. Mice were monitored for survival (A), weight loss (B), and disease score (as described in Methods) (C) for 12 days post-infection (dpi). WT, n=4 mice; IA, n=8 mice; FA, n=8 mice. (D) Brains were collected at 5 dpi and titers were determined by plaque assay. WT, n=6; IA, n=7; FA, n=8. The data show the combined results from two independent experiments.

### MERS-CoV I1238A and F1239A proteins have opposing effects on Mac1 biochemical activities

To determine how these mutations impact the biochemical activities of Mac1, we aimed to purify Mac1 protein with these mutations and utilize *in vitro* assays to measure ADP-ribosylhydrolase and ADP-ribose binding activity of each mutant protein. Multiple attempts to produce MHV Mac1 protein failed, so we engineered these mutations into the *Merbecovirus* MERS-CoV and the *Sarbecovirus* SARS-CoV-2 Mac1 recombinant proteins, as we’ve previously produced WT Mac1 proteins from each virus (7). We first produced soluble I1238A and F1239A MERS-CoV Mac1 proteins and performed isothermal titration calorimetry (ITC) to measure Mac1-ADP-ribose binding. ITC measures the release or absorption of energy during a binding reaction and has been used extensively to measure macrodomain-ADP-ribose interactions (7, 23, 27, 36, 37). Compared to WT protein, the F1239A protein bound to free ADP-ribose with a substantially higher K_D_ value (60 µM vs. 7.2 µM) indicating reduced binding ability. In contrast, the I1238A protein bound to ADP-ribose with a K_D_ nearly equivalent to that of WT (12.7 µM vs 7.2 µM) (Fig. 4A). In addition to ITC, we also performed an AlphaScreen assay, as previously described (38–40), to determine the ability of each protein to bind to a non-cleavable ADP-ribosylated peptide. This peptide was chosen considering the robust signal that it produces in the presence of Mac1 proteins (38). Similar to the ITC assay, the F1239A Mac1 had reduced AlphaScreen counts at all concentrations of protein tested as compared to WT protein, indicating poor binding to the ADP-ribosylated peptide (Fig. 4B). Remarkably, the I1238A Mac1 protein had substantially increased AlphaScreen counts at all concentrations of protein tested, indicating that this mutation has enhanced binding to an ADP-ribosylated peptide (Fig. 4B). As a control, we found that the I1238A protein did not bind to the unmodified peptide (data not shown). To further test this observation, we performed a competition assay by adding increasing amounts of ADP-ribose to the reaction. ADP-ribose inhibited the peptide-ADP-ribose interaction of WT MERS-CoV protein with an average IC_50_ value, of 0.155 *μ*M, while it had a much higher IC_50_ value of 1.6 *μ*M for the I1238A protein. These results demonstrate that I1238A had a stronger interaction with an ADP-ribosylated peptide, but not free ADP-ribose, than WT protein (Fig. 4C).

**Figure 4.**
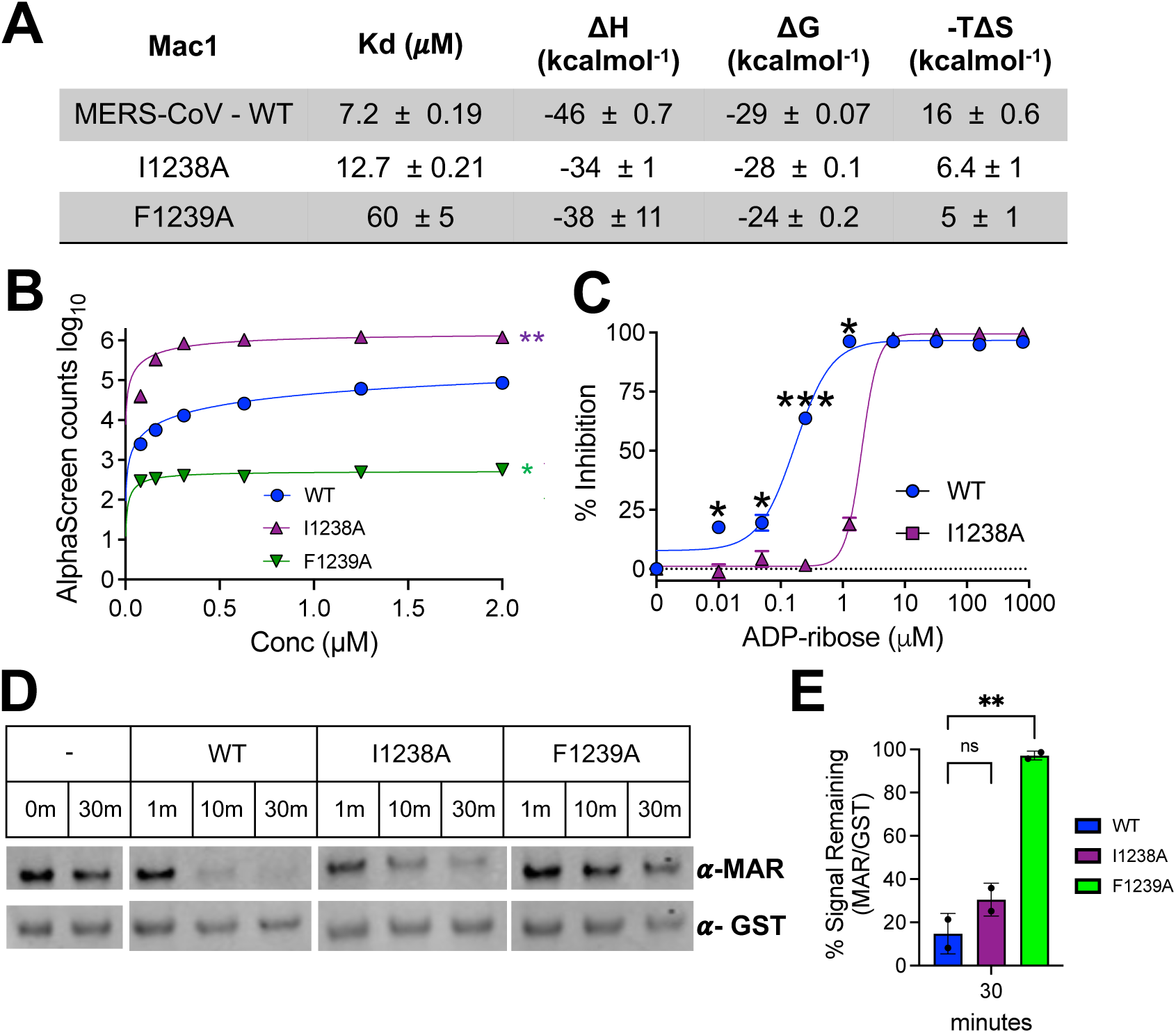
MERS-CoV I238A and F1239A Mac1 mutations have opposing effects on ADP-ribose binding and hydrolysis. (A) Mac1 protein was incubated with free ADP-ribose and binding affinity was measured by isothermal calorimetry as described in Methods. (B) An ADP-ribosylated peptide was incubated with indicated macrodomains at increasing concentrations and Alphacounts were measured as described in Methods. (C) ADP-ribose (ADPr) competition assays were used to block the interaction between macrodomain proteins and ADP-ribosylation peptides in the AS assay. Data was analyzed as described in Methods. The data in A-C represent combined results of 2 independent experiments for each protein. (D) WT, I1153A, and F1154A MERS-CoV Mac1 proteins were incubated with MARylated PARP10 CD *in vitro* at an [E]/[S] molar ratio of 1:5 for the indicated times at 37°C. ADP-ribosylated PARP10 CD was detected by IB with anti-ADP-ribose binding reagent (MAB1076; MilliporeSigma) while total PARP10 CD protein levels was detected by IB with GST antibody. The reaction with PARP10 CD incubated alone at 37°C was stopped at 0 or 30 min. The image in D is representative of 2 independent experiments. (E) The level of de-MARylation was measured after 30 minutes by quantifying relative band intensity (ADP-ribose/GST-PARP10) using ImageJ software. Error bars represent standard deviations. The results in E are the combined resulted of 2 independent experiments.

We next tested the ability of the MERS-CoV I1238A and F1238A Mac1 proteins to hydrolyze mono-ADP-ribose (MAR) from protein as previously described (7). The WT, I1238A, and F1239A Mac1 proteins were incubated with MARylated PARP10 at a 1:5 enzyme to substrate ([E]/[S]) ratio, and the reaction was stopped at several timepoints to determine the ability of each protein to hydrolyze MAR. As a control, MARylated PARP10 was collected at the first (0 min) and the final (30 min) timepoints. Over the course of 30 minutes, the MERS-CoV I1238A Mac1 protein decreased the level of MARylated PARP10 to levels similar to that of the MERS-CoV WT Mac1 protein, while the MERS-CoV F1239A Mac1 protein did not efficiently remove MAR from PARP10 (Fig. 4D-E). Taken together, we conclude that the MERS-CoV I1238A and F1239A mutations had somewhat opposing effects on the activity of Mac1. While F1239A mutant Mac1 protein has decreased ADP-ribose binding and hydrolysis activity, the I1238A Mac1 has at least similar, if not increased ADP-ribose binding, depending on the substrate, with only a modest reduction in enzyme activity compared to the MERS-CoV WT Mac1 (Table 1).

**Table 1.** Summary of Mac1 ADP-ribose binding, hydrolysis, and replication activity.

| **Virus** |  | **MERS-CoV** |  |  | **SARS-CoV-2** |  |  |  |
| --- | --- | --- | --- | --- | --- | --- | --- | --- |
| **Mutation** | WT | N1147A | I1238A** | F1239A | WT | N1062A* | I1153A | F1154A |
| **Binding** | + | nd | +/++ | - | + | + | ++ | ++ |
| **Hydrolysis\*** | + | nd | + | - | + | - | + | - |
| **Replication\*\*\*** | + | - | - | - | + | - | - | - |

nd, not determined
(+) - WT levels
(++) - significantly increased compared to WT
(-) - significantly decreased compared to WT
\*SARS-CoV-2 N1062A binding and hydrolysis was not directly compared with I1153A and F1154A
\*\* MERS-CoV I1238A binding results depend on the assay
\*\*\*SARS-CoV-2 replication is defined in the presence of IFNγ

### MERS-CoV I1238A and F1239A viruses have decreased virus production in Calu-3 cells

We next tested the ability of MERS-CoV N1147A, I1238A, and F1239A recombinant viruses to replicate in cell culture. First, using recombination, we inserted a GFP cassette in place of ORF5 in the MERS-CoV-MA BAC, as ORF5 quickly mutates in cell culture, which could complicate our results (41, 42). Considering that the I1238A had equivalent or enhanced biochemical activities compared to WT protein, we hypothesized that only the N1147A and F1239A viruses would impact MERS-CoV replication, similar to results seen with MHV (Figs. 2-3). Each mutant virus produced infectious virus near WT levels at 24 and 48 hpi in Vero81 cells, which are unable to produce interferon (IFN) (Fig. 5A). Next, we tested the replication of these viruses in IFN-competent Calu-3 cells, which are human bronchial epithelial cells that are a commonly used cell line for both MERS-CoV and SARS-CoV-2 infections. To our surprise, all 3 viruses replicated poorly in these cells. In Calu3 cells each virus had between 2.1 and 2.5-fold lower levels of infectious virus produced than WT virus at 48 hpi, and had between 8.5 and 10.5-fold lower levels of infectious virus production at 72 hpi (Fig. 5B). As each mutation led to a nearly identical reduction in virus replication, we conclude that each of these residues is critical for MERS-CoV replication in cell culture, and that the defect of the I1238A virus could be due to enhanced ADP-ribose binding (Table 1).

**Figure 5.**
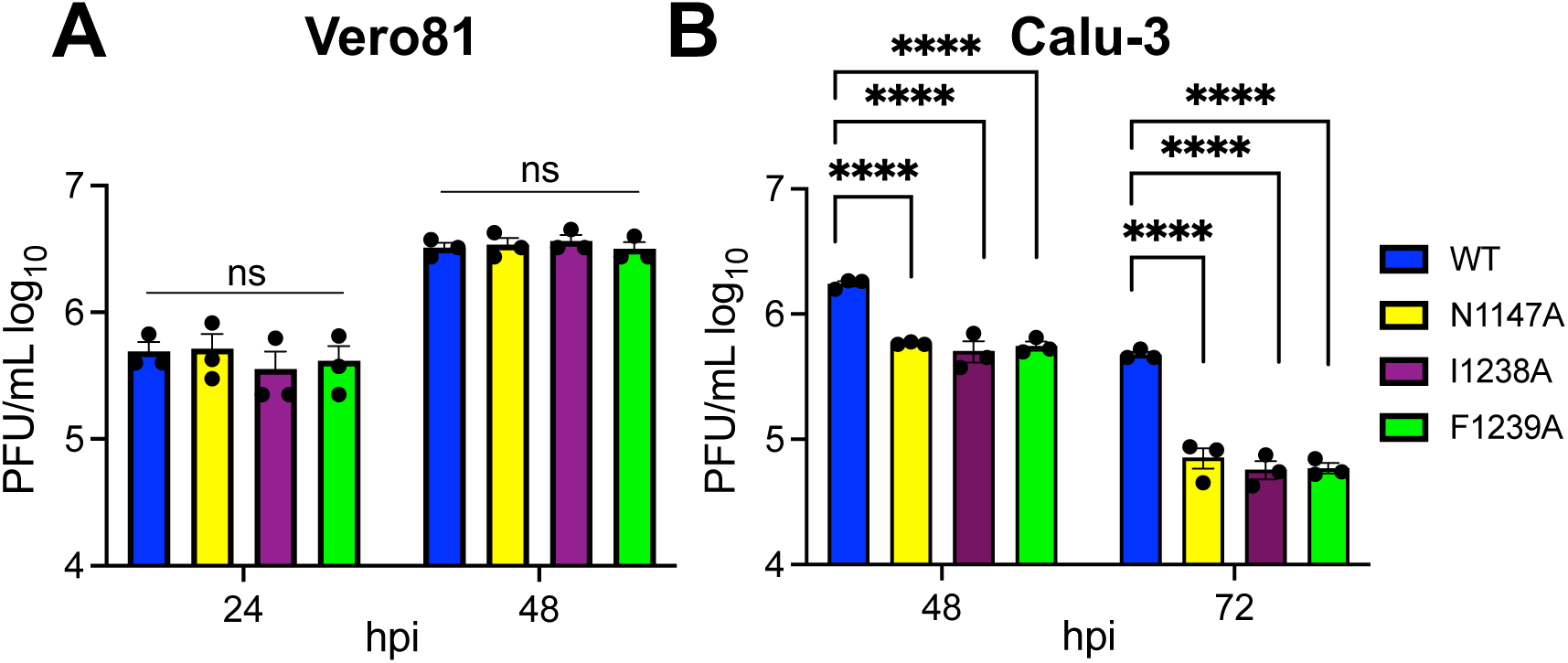
MERS-CoV I1238A and F1239A have similarly decreased replication in Calu-3 cells. (A-B) Vero81 (A) and Calu3 (B) cells were infected at an MOI of 0.1 PFU/cell. Cells and supernatants were collected at indicated times post-infection (hpi) and progeny virus was measured by plaque assay. The data in panels A and B show one experiment representative of two (A) or three (B) independent experiments with n = 3 for each experiment.

### SARS-CoV-2 I1153A and F1154A have increased ADP-ribose binding activity

The MERS-CoV data indicated that increased ADP-ribose binding activity may lead to replication defects in culture. However, the I1238A MERS-CoV mutant that had increased ADP-ribose binding, had at least a modest defect in enzyme activity, which could account for its poor replication (Table 1). To further test the hypothesis that increased ADP-ribose binding could be detrimental to infection, we engineered these mutations in SARS-CoV-2 to analyze their impact on Mac1 biochemical functions and viral replication. We produced soluble I1153A and the F1154A SARS-CoV-2 Mac1 proteins and first performed ITC to determine the ADP-ribose binding ability of each Mac1 mutant protein. Interestingly, both the SARS-CoV-2 I1153A and the F1154A Mac1 proteins had increased binding to free ADP-ribose, with K_D_ values of 5.49 µM and 5.11 µM, respectively, compared to the K_D_ value of 16.8 µM for WT protein (Fig. 6A). To confirm these results with free ADP-ribose, we performed a differential scanning fluorimetry assay, which measures the thermal stability of a protein in the presence of a substrate. Again, both the I1153A and F1154A proteins had increased binding as compared to WT protein in this assay (Fig. 6B). Finally, we tested the ability of each protein to bind to the ADP-ribosylated peptide in the AlphaScreen assay. Unfortunately, the F1154A, but not the I1153A protein strongly bound to the control, unmodified peptide (Fig. 6C). Thus, we were unable to evaluate the ability of F1154A protein’s ability to bind ADP-ribosylated peptide. However, the I1153A protein had significantly increased binding to the ADP-ribosylated peptide compared to the WT Mac1 protein (Fig. 6D). We conclude that both the I1153A and F1154A proteins have increased binding to ADP-ribose *in vitro*.

**Figure 6.**
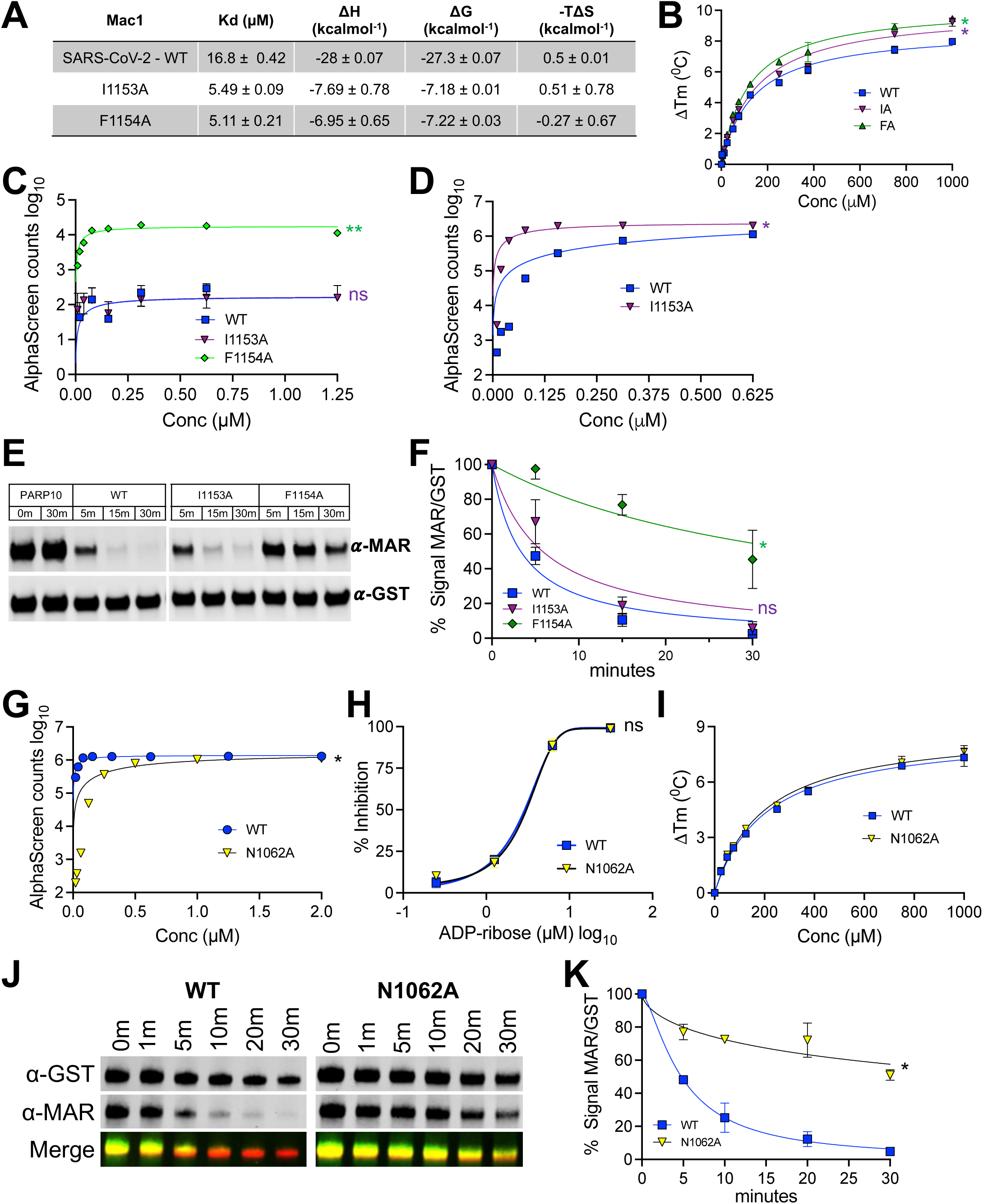
SARS-CoV-2 Mac1 mutations impact both ADP-ribose binding and hydrolysis. (A) SARS-CoV-2 Mac1 protein was incubated with free ADP-ribose and binding affinity was measured by isothermal calorimetry as described in Methods. (B) WT, I1153A, and F1154A Mac1 proteins (10 μM) were incubated with increasing concentrations of ADP-ribose and measured by DSF as described in Methods. The data in panels A and B show one experiment representative of three independent experiments. (C) A control (non-ADP-ribosylated) peptide was incubated with indicated Mac1 proteins at increasing concentrations and Alphacounts were measured as described in Methods. (D) An ADP-ribosylated peptide (same peptide sequence as in (C)) was incubated with indicated Mac1 proteins at increasing concentrations and Alphacounts were measured as described in Methods. The data in panels C and D show one experiment representative of two independent experiments. (E) WT, I1153A, and F1154A SARS-CoV-2 Mac1 proteins were incubated with MARylated PARP10 CD *in vitro* at an [E]/[S] molar ratio of 1:5 for the indicated times at 37°C. ADP-ribosylated PARP10 CD was detected by IB with anti-ADP-ribose binding reagent (MAB1076; MilliporeSigma) while total PARP10 CD protein levels were detected by IB with GST antibody. The reaction with PARP10 CD incubated alone at 37°C was stopped at 0 or 30 min. The data in panels D shows one experiment representative of three independent experiments. (F) The level of de-MARylation from 3 separate experiments was measured by quantifying relative band intensity (ADP-ribose/GST-PARP10) using ImageJ software. Intensity values were plotted and fitted to a nonlinear regression curve; error bars represent standard deviations. (G) ADP-ribosylated peptide was incubated with WT and N1062A Mac1 proteins at increasing concentrations and Alphacounts were measured as described in Methods. (H) ADP-ribose (ADPr) competition assays were used to block the interaction between WT and N1062A Mac1 proteins and ADP-ribosylated peptides. Data was analyzed as described in Methods. The data represent the means ± SD of 2 independent experiments for each protein. (I) WT and N1062A Mac1 proteins (10 μM) were incubated with increasing concentrations of ADP-ribose and measured by DSF as described in Methods. (J) WT and N1062A SARS-CoV-2 Mac1 proteins were incubated with MARylated PARP10 CD *in vitro* at an [E]/[S] molar ratio of 1:5 for the indicated times at 37°C. ADP-ribosylated PARP10 CD was detected by IB with anti-ADP-ribose binding reagent (green) while total PARP10 CD protein levels were detected by IB with GST antibody (red). The reaction with PARP10 CD incubated alone at 37°C was stopped at 0 or 30 min. The data is representative of 2 independent experiments. (K) The level of de-MARylation in D was measured by quantifying relative band intensity from 2 independent experiments (ADP-ribose/GST-PARP10) using ImageJ software. Intensity values were plotted and fitted to a nonlinear regression curve. The data represent the means ± SD of 2 independent experiments for each protein.

Next, we tested the ability of each SARS-CoV-2 protein to remove MAR from MARylated PARP10, again at a 1:5 [E]/[S] ratio to account for defects in enzyme turnover. Like MERS-CoV F1239A, SARS-CoV-2 F1154A had only modest hydrolysis activity. In contrast, I1153A Mac1 protein had robust enzymatic activity, which was virtually indistinguishable from WT protein (Fig. 6E-F), which is consistent with previously published results (9). These results demonstrate that the I1153A and F1154A both have enhanced ADP-ribose binding, but that only F1154A has reduced enzymatic activity (Table 1).

As many previous studies on the role of Mac1 in virus replication include the highly conserved asparagine-to-alanine mutation, we also generated an N1062A SARS-CoV-2 Mac1 protein. While CoVs with this mutation are easily recovered and often grow like WT virus, unlike full deletion viruses (24, 34), for unknown reasons this recombinant protein is highly unstable. Thus, several modifications to the normal protocol were made to create a small amount of soluble protein. While the small amount of protein did not allow for ITC measurements, this protein had similar ADP-ribose binding properties as WT protein, as determined in combination by the alphascreen, ADP-ribose competition, and differential scanning fluorimetry assays (Fig. 6G-I). In contrast, this protein had substantially reduced ADP-ribosylhydrolase activity (Fig. 6J-K), indicating that this mutation primarily impacts the enzyme activity of Mac1. Previously published data also supports the hypothesis that this mutation primarily impacts the enzyme activity of Mac1 (11, 12, 23, 29, 31, 36). Thus, a combination of recombinant N1062A, I1153A, and F1154A SARS-CoV-2 viruses may be useful in defining the role of ADP-ribose binding and hydrolysis during infection, as N1062A has similar binding and reduced enzyme activity, F1154A has increased binding and reduced enzyme activity, and I1153A has only increased binding (Table 1).

### I1153A and F1154A mutations increase the sensitivity of SARS-CoV-2 to cIFNγ

We previously reported that a Mac1 deleted SARS-CoV-2 replicates at levels similar to WT virus, but was highly sensitive to IFNγ, but not IFN*β* pretreatment in Calu-3 cells (24). We next tested the ability of I1153A and F1154A recombinant viruses to produce infectious virus in the presence and absence of IFNγ in both A549-ACE2 and Calu-3 cells. Without IFNγ pre-treatment at 48 hpi, both I1153A and F1154A produce infectious virus at WT levels, as expected (Fig. 8A). In contrast, there is a substantial decrease in the production of both I1153A and F1154A infectious virus compared to WT SARS-CoV-2 in the presence of IFNγ in both cell lines (Fig. 8A-B). Furthermore, mutation at the N1062 residue has increased sensitivity to IFNγ as well, though not as severe as the I1153A and F1154A mutant viruses (24). We conclude that the I1153A and F1154A mutations are detrimental for the ability of SARS-CoV-2 to replicate efficiently in the presence of IFNγ.

### SARS-CoV-2 I1153A and F1154A are attenuated in K18-ACE2 mice

Next, we tested whether the I1153A and F1154A Mac1 mutations would be detrimental to SARS-CoV-2 infection in mice. Previously, a SARS-CoV-2 Mac1 deletion virus was shown to be extremely attenuated in K18-ACE2 mice, while the N1062A mutant was mildly attenuated, with approximately 50% of mice surviving the infection (24, 29). We hypothesized that like the SARS-CoV-2 N1062A mutant, there would be at least partial attenuation of I1153A and F1154A viruses in mice. Following an intranasal infection, both the SARS-CoV-2 I1153A and F1154A viruses were extremely attenuated in mice, as they did not cause any weight loss or lethal disease in mice, similar to *Δ*Mac1, whereas WT SARS-CoV-2 causes 100% mortality by 9 dpi (Fig. 9A-B). Viral titers were reduced by ∼4-5 fold at 1 dpi (Fig. 9C), and by 8 dpi both the I1153A and F1154A viruses were cleared from the lungs of mice (Fig. 9D). Furthermore, mice infected with these viruses had reduced signs of disease, such as bronchointerstitial pneumonia, edema, or fibrin, as measured by H&E staining (Fig. 9E-F). Finally, both I1153A and F1154A infected mice had significantly increased levels of IFN-I, IFN-III, ISG15, and CXCL-10 mRNA, similar to *Δ*Mac1 infection levels (Fig. 9G) (29, 34). These results demonstrate that both I1153A and F1154A mutations are detrimental to SARS-CoV-2 replication and pathogenesis *in vivo*.

In total, while the highly conserved isoleucine and phenylalanine mutations in MERS-CoV and SARS-CoV-2 have different effects on Mac1 biochemical activities *in vitro*, their impact on virus replication and pathogenesis were remarkably similar (Table 1). The simplest explanation is that enhanced ADP-ribose binding has a detrimental effect on Mac1’s function, and that the conserved isoleucine residue acts to control ADP-ribose binding to allow for optimal function. But how does this isoleucine residue control ADP-ribose binding? Prior NMR data from the Venezuelan equine encephalitis virus (VEEV) macrodomain indicated that prior to ADP-ribose binding there is a significant transition that increases the distance between loop 1 and loop 2 from 7 to 10 Å to accommodate ADP-ribose as a substrate (43). We hypothesized that the I1153A protein has increased binding because the protein no longer requires this transition to bind ADP-ribose due to the loss of the bulky isoleucine side chain. To support this hypothesis, we performed a molecular dynamic (MD) simulation of the I1153A and WT proteins in the presence and absence of ADP-ribose and measured the 1 ns running average distance between the I or A 1153 residue and G1069 (Fig. 10A-C). In the absence of ADP-ribose, the mutant protein (A1153) consistently sampled conformations containing a larger distance between these residues, around or longer than ∼7.5 Å, which results in a largely open crevice between the two loops (Fig. 10A). In contrast, the distance between these residues for the WT protein (I1153) was more often less, even sampling distances below 5 Å (Fig. 10A), and the crevice appears mostly in a closed state, only occasionally opening wide enough to allow for ADP-ribose binding (Fig. 10B-C). However, in the presence of ADP-ribose, these residues were nearly the same distance apart in WT and I1153A Mac1 proteins, ∼7.5-8 Å, throughout the simulation, indicating that the mutation may not impact the ability of ADP-ribose to exit the binding site. These results suggest that the isoleucine , due to its bulky side-chain, controls the ability of ADP-ribose to enter, but not exit, the ADP-ribose binding domain.

## DISCUSSION

Research on the CoV Mac1 domain over the last two decades has established this protein domain as critical for the replication and pathogenesis of CoVs (44). However, the relative contributions of its two major biochemical activities, ADP-ribose binding and deMARylation, to its function during infection has remained unclear. Most research on the CoV Mac1 domain has utilized the mutation of a highly-conserved Asn-to-Ala to understand its role in viral replication and pathogenesis. This mutation nearly eliminates the ADP-ribosylhydrolase activity of SARS-CoV Mac1, however, it was previously not clear how much this mutation impacts ADP-ribose binding. Results from archaeal and alphavirus macrodomains have indicated that mutation of the orthologous asparagine residue in those proteins to alanine modestly reduces ADP-ribose binding (23, 36). Here we created an N1062A Mac1 protein from SARS-CoV-2 and found that it had a severe defect in enzyme activity, but only had a modest, but significant reduction in ADP-ribose binding compared to WT protein (Fig. 7). This confirms that this residue plays a large role in ADP-ribosylhydrolase activity but only moderately impacts ADP-ribose binding. These results further indicate that phenotypes associated with this mutation, including increased IFN production and enhanced sensitivity to IFN*β* and IFN*γ*, are likely due to the loss of ADP-ribosylhydrolase activity, especially considering a SARS-CoV-2 aspartic acid to alanine mutation (D1044A), predicted to dramatically reduce binding, was not sensitive to IFNγ and the orthologous MHV mutation, D1329A does not substantially increase IFN*β* production (7, 34).

**Figure 7.**
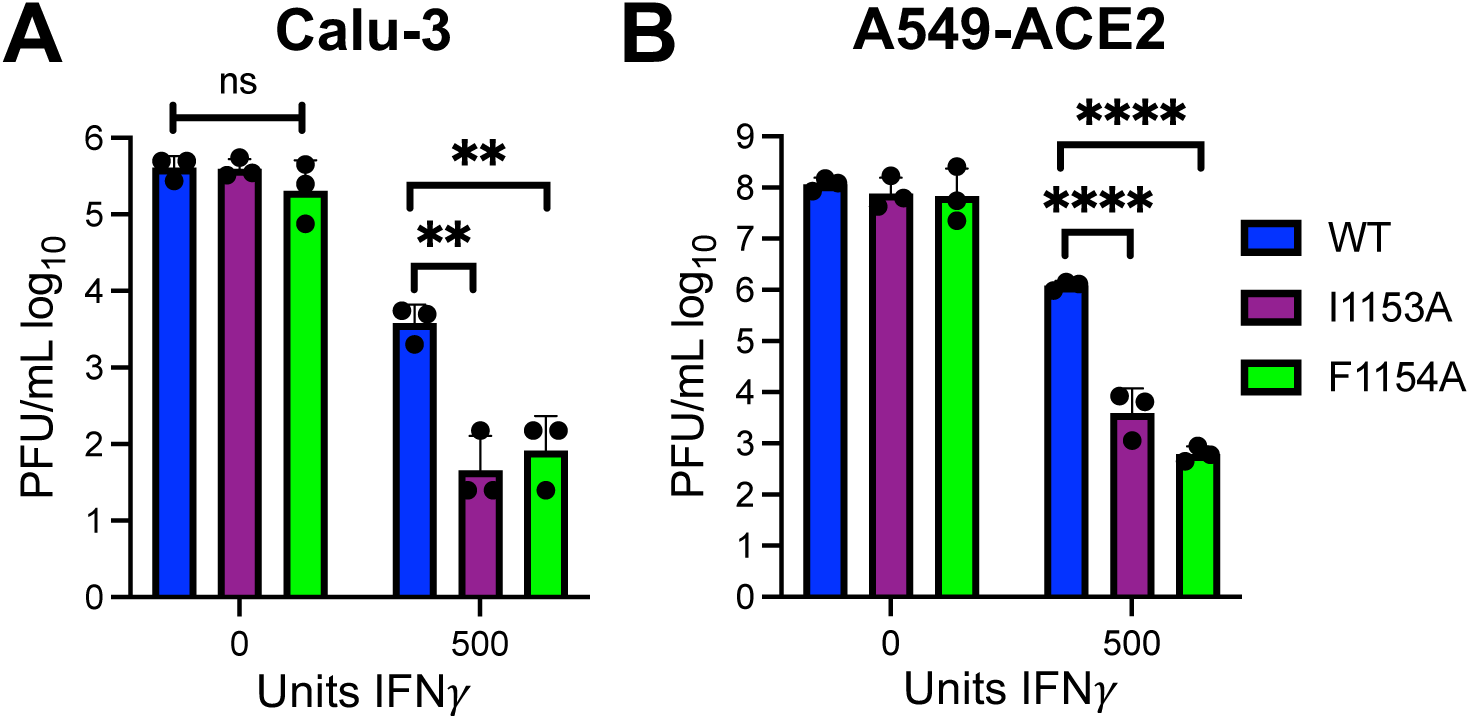
SARS-CoV-2 Mac1 I1153A and F1154A mutations have detrimental effects on SARS-CoV-2 replication in the presence of IFNγ. Calu-3 (A) and A549-ACE2 (B) cells were pretreated with 500 units of IFNγ for 18-20 hours prior to infection. Then cells were infected at an MOI of 0.1 PFU/cell. Cells and supernatants were collected at 48 hpi and progeny virus was measured by plaque assay. The data in panels A and B show one experiment representative of three independent experiments with *n* = 3 for each experiment.

**Figure 8.**
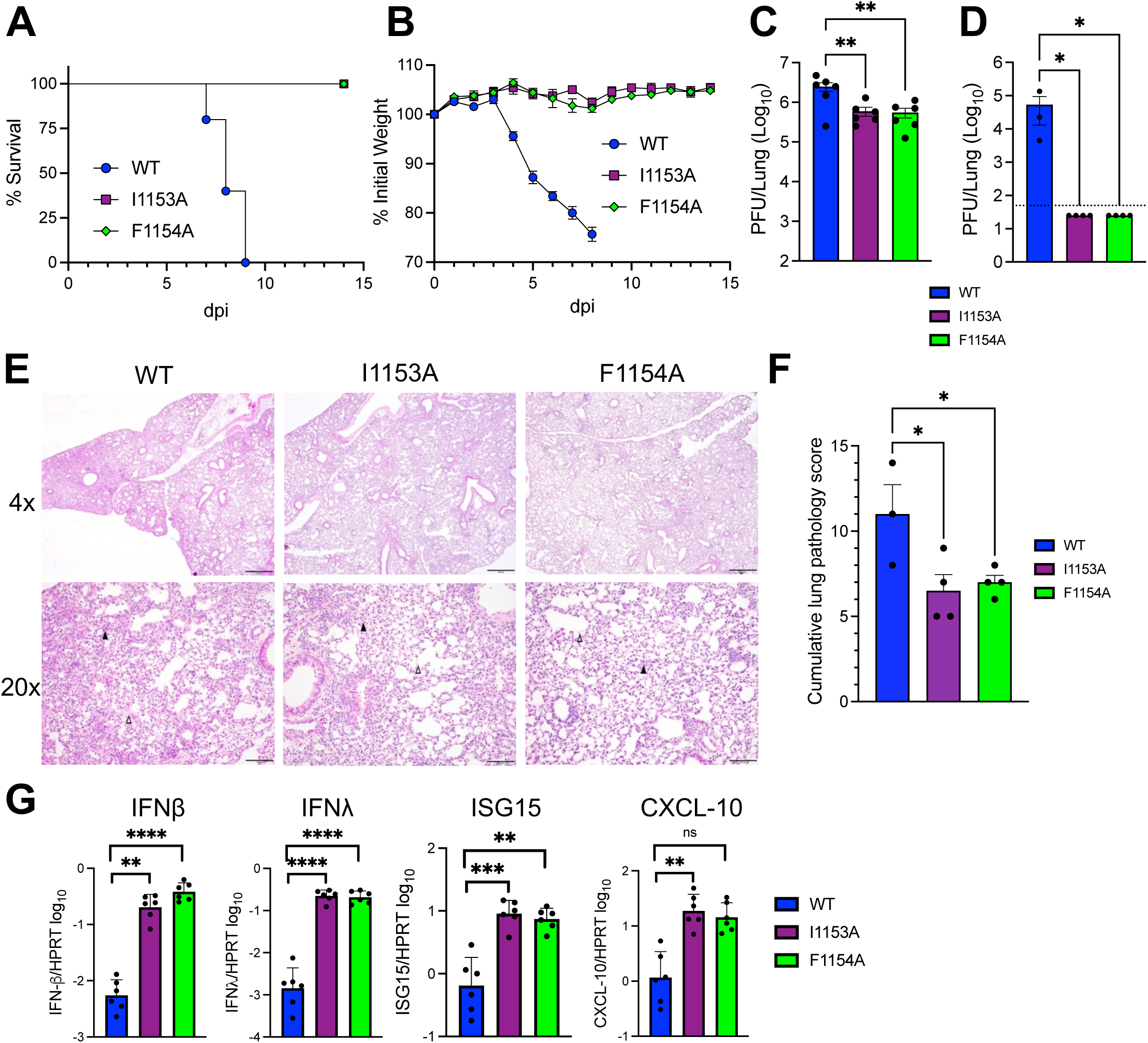
SARS-CoV-2 I1153A and F1154A are highly attenuated and induce elevated innate immune responses in the lungs of infected mice. K18-ACE2 C57BL/6 mice were infected i.n. with 2.5 × 10^4^ PFU of virus. (A-B) Survival (A) and weight loss (B) were monitored for 14 days. n=5 for survival and n=9 for weightloss for all groups. (C) Lungs were harvested at 1 dpi and viral titers were determined by plaque assay. n=6 for all groups. (D) Lungs were harvested at 8 dpi and viral titers were determined by plaque assay. Dotted line indicates limit of detection. n=3 for WT, n=4 for I1153A and F1154A. (E) Photomicrographs (hematoxylin and eosin stain) of lungs infected mice at 8 dpi demonstrating bronchointerstitial pneumonia (black arrow) and edema and fibrin (open arrow). (F) Mice were scored for bronchointerstitial pneumonia, inflammation, and edema/fibrin deposition (each on a 0-5 scale). Bar graphs represent cumulative lung pathology score in WT n=3, I1153A n=4, F1154A n=4. (G) Lungs were harvested at 1 dpi in Trizol and RNA was isolated. Transcripts levels were determined using qPCR with the ΔCT method. n=6 for all groups.

**Figure 9.**
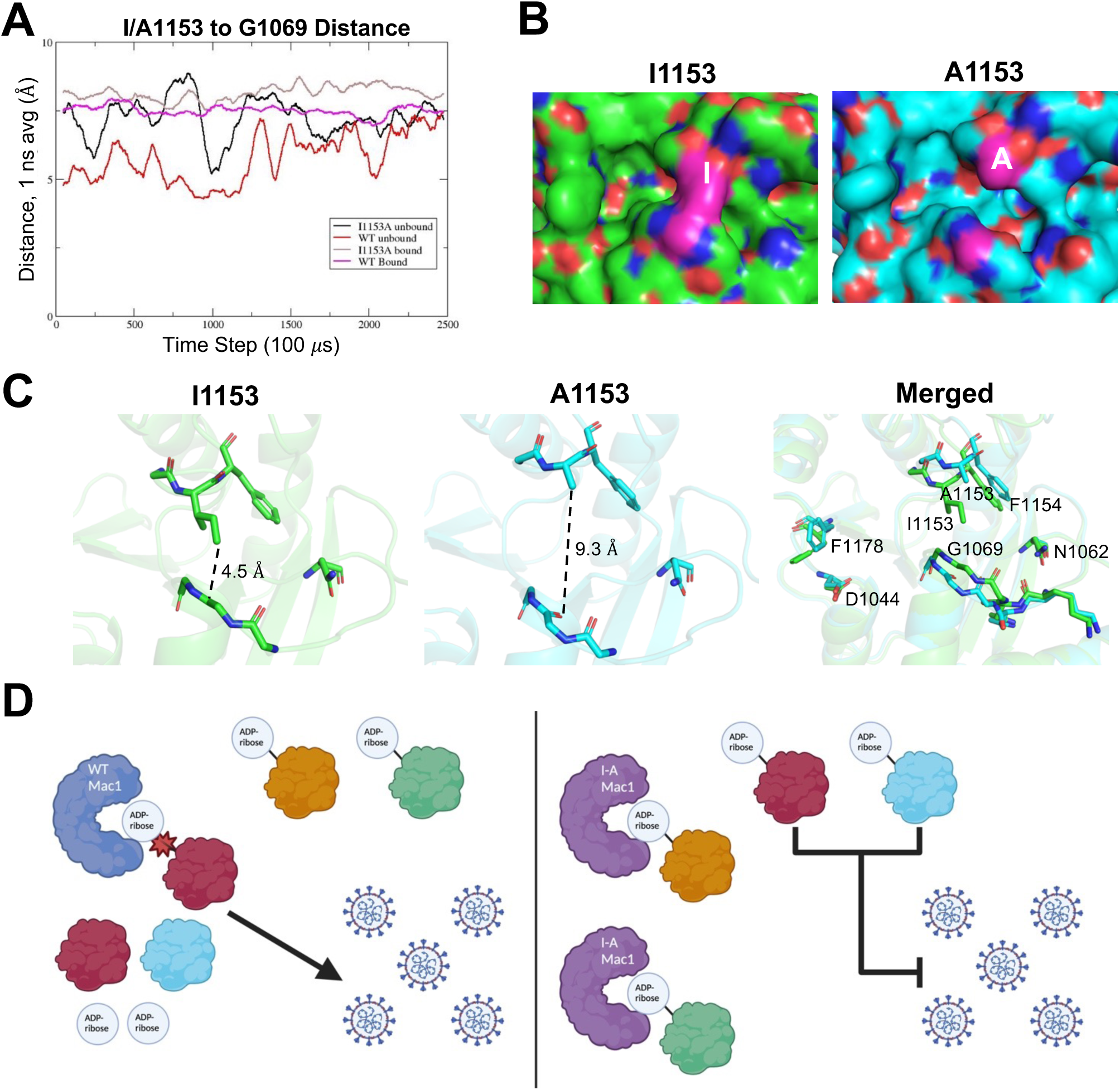
Models of isoleucine-to-alanine mutation on Mac1 structure and virus replication. (A) Molecular simulation of the ADP-ribose binding domain of the SARS-CoV-2 Mac1 protein was performed in absence and presence of ADP-ribose. The 1 ns averaged I/A1153 to G1069 distance was measured through the course of four 25 ns MD simulations of ADP-ribose bound and unbound WT and I1153A protein. (B-C) A representative image at 12 ns of the simulation demonstrating the distance between the I1153 and A1153 residues and G1069 at 12 ns into the simulation without ADP-ribose in a space-filling (B) or stick model (C). (D) (Left) In the presence of IFNγ, the WT SARS-CoV-2 Mac1 removes ADP-ribose from specific proteins (red and blue) that have an ADP-ribose on an acidic reside which enhances virus replication. (Right) Due to the open conformation of SARS-CoV-2 I1153A Mac1 protein, it binds to ADP-ribose bound to non-acidic residues (gold and green). Since Mac1 cannot remove proteins from non-acidic residues, this limits its ability to interact with relevant substrate, and the ADP-ribose remains on its normal target proteins leading to poor virus replication.

This study focused on the isoleucine and phenylalanine residues located in loop 2 of Mac1, near this asparagine residue, and their roles in the biochemical activites of Mac1 and how they impact CoV replication and pathogenesis. The isoleucine in loop 2 of the CoV Mac1 protein has been described as a bridge that extends from loop 2 to loop 1 that covers the phosphate binding domain of Mac1, forming a narrow channel that might impact binding or hydrolysis (7, 10, 11, 35). Furthermore, this residue participates in the transition of these loops from the apo form to the ADP-ribose bound form (43), again indicating that this residue may impact ADP-ribose binding. Somewhat surprisingly, we found that an I-A mutation instead led to enhanced ADP-ribose binding based on a peptide-ADP-ribose binding assay for both the MERS-CoV and SARS-CoV-2 Mac1 proteins (Figs. 4B-C, 6A-B). Modeling data indicates that with this mutation, the distance between the two loops is consistently large enough that Mac1 can likely accept substrates more readily, as opposed to Mac1 with the isoleucine (Fig. 10A-C). Following ADP-ribose binding, the I-A mutation does not appear to impact the distance between the loops, perhaps explaining why the hydrolysis activity of Mac1 was not affected for either Mac1 protein. These results suggest that the isoleucine residue serves as a gate to control ADP-ribose binding levels.

In contrast, the phenylalanine residue forms van der Waals interactions with the distal ribose, and similar to the nearby asparagine residue, appears to help position the ribose for hydrolysis. Biochemical data has supported those predictions, as mutations of this residue generally result in substantial loss of hydrolysis activity, which we observed here for both the MERS-CoV and SARS-CoV-2 Mac1 proteins (Figs. 4D-E, 6C-D). Interestingly, the F-A mutation had diverse roles in ADP-ribose binding. For MERS-CoV Mac1, this mutation led to reduced binding, while for SARS-CoV-2 this mutation enhanced both free and peptide-conjugated ADP-ribose binding. As the phenylalanine residue resides just outside the terminal ribose, it’s conceivable that in some cases this residue may occlude ADP-ribose binding during its transitions, while in others it may be just far enough away to not impact the ability of ADP-ribose to enter the binding pocket (7, 43) Furthermore, an orthologous tyrosine to alanine mutation in the CHIKV macrodomain, Y114A, also led to enhanced ADP-ribose binding, indicating that this residue helps control ADP-ribose binding in multiple macrodomains. While it’s unclear how these identical mutations in the MERS-CoV and SARS-CoV-2 Mac1 proteins had opposing effects on ADP-ribose binding, it highlights the difficulty in attributing specific biochemical roles for individual residues from one macrodomain to another. In addition, there are several limitations to our approaches. As there are no known substrates of Mac1 in infection, we use artificial substrates for our *in vitro* assays. These include free ADP-ribose, the product of Mac1’s hydrolytic function, an ADP-ribosylated peptide using a random peptide sequence, and an auto-MARylated PARP10 protein. However, these are standard assays in the field (9, 23, 27, 35, 45, 46). Furthermore, our modeling data is based on molecular dynamic simulations approximated using the crystal structure of the SARS-CoV-2 Mac1 protein, as no crystal structure exists for the I1153A mutation, and no NMR data is available for either protein. Finally, free ADP-ribose produced during the de-ADP-ribosylation assays could impact the assay, though our deADP-ribosylation experiments performed at a low [E]/[S] ratio suggests that this is not the case (Figs. 4, 6).

Both MERS-CoV and SARS-CoV-2 I-A and F-A mutations were equally attenuated in both cell culture and in mice despite having somewhat distinct biochemical properties (Figs. 5, 7, 8). The MERS-CoV mutant viruses replicated normally in Vero81 cells but replicated poorly in Calu-3 cells at levels similar to the N1147A virus. Furthermore, both F-A and I-A SARS-CoV-2 mutant viruses replicated poorly following IFN-*γ* treatment and induced high levels of IFN and ISG levels following infection in mice, again demonstrating that despite having unique biochemical properties, these mutations led to very similar virological phenotypes (Figs. 7, 9). There are several potential hypotheses that might explain this, and we will outline some of the possibilities here. Our first hypothesis is that increasing ADP-ribose binding could lead to decreased enzyme activity during infection. Why might an increase in ADP-ribose binding lead to reduced enzyme activity during infection? One possibility is that enhanced binding would negatively affect enzyme turnover. However, our *in vitro* enzyme assays were performed at an [E]/[S] ratio of 1:5, indicating that the mutant protein has largely normal enzyme turnover. ADP-ribose can be covalently attached to several different amino acids, including cysteine, serine, arginine, glutamic and aspartic acid, but the MacroD2 class of macrodomains primarily removes ADP-ribose from acidic residues. Therefore, a second hypothesis is that enhancing the ADP-ribose binding abilities of Mac1 may cause it to bind to proteins with ADP-ribose attached at non-acidic residues that it can’t remove and are not relevant for virus infection. For instance, during infection WT Mac1 primarily engages with either anti- or pro-viral proteins that are MARylated on an acidic residue. Mac1 removes these modifications, which promotes virus replication and pathogenesis. In contrast, Mac1 I-A may bind non-specifically to proteins MARylated at non-acidic residues, such as serine or asparagine, reducing its ability to engage with its primary targets. In this case, Mac1 becomes stuck to irrelevant targets, while its main target proteins remain ADP-ribosylated, leading to reduced virus replication (Fig. 10D). However, this is not the only possible hypothesis. Additional hypotheses include but are not limited to: *i)* Enhanced ADP-ribose binding enables Mac1 to bind with additional ADP-ribosylated protein or RNA substrates, which is detrimental to infection; *ii)* the I-A mutation could enhance the binding to non-ADP-ribosylated proteins, such as PLPro, which was previously shown to bind the MHV Mac1 protein (47); and *iii)* the I-A mutation could either increase, or decrease the enzyme activity of Mac1 for alternative substrates, such as NAD^+^, ADP-ribose-1’-phosphatase, or O-acetyl-ADP-ribose (11, 31, 48). CoV infection can dysregulate the NAD^+^ system and reduce NAD^+^ levels. We found that NAD^+^ levels are reduced following MHV infection and that this depletion enhances MHV Mac1 mutant virus replication, while others have shown that NAD^+^ enhancing treatments provide mice some level of increased protection from SARS-CoV-2 (49–51). However, macrodomains are not known to be major consumers of NAD^+^, thus this possibility seems unlikely. Furthermore, there’s no indication that ADP-ribose-1’-phosphatase, or O-acetyl-ADP-ribose impact CoV infection. Additional experiments will need to be designed to demonstrate how the I-A mutation results in attenuated virus replication.

The function of the isoleucine residue of the MHV Mac1 protein appears to be unique, as the mutation of I-A had little-to-no impact on virus replication. As we have been unable to purify the WT MHV Mac1 protein in bacteria, we can only speculate as to how this mutation impacts ADP-ribose binding and hydrolysis. The simplest hypothesis is that this mutation does not enhance ADP-ribose binding as it did for MERS-CoV or SARS-CoV-2. Alternatively, as MHV appears to be highly dependent on the ADP-ribose binding function of Mac1 (24, 34), an increase in ADP-ribose binding may have some beneficial outcome that counteracts the negative effects of this mutation, resulting in a virus that replicates much like WT. Conversely, the F1441A mutant virus replicates poorly in all cells tested and was attenuated in mice, similar to a D1329A mutant virus, a mutation predicted to largely impact ADP-ribose binding. In addition, it was partially, but not fully, rescued in PARP12 KO cells, which we previously found rescued N1347A, but had no effect on D1329A. Thus, based on these and prior results with N1347A and D1329A, we hypothesize that this mutation reduces both enzyme and binding activity, similar to the MERS-CoV F1239A Mac1 protein.

These results provide new insights into how Mac1 regulates ADP-ribose binding for its benefit, which could have important implications for the development of inhibitors targeting Mac1. Finally, these mutations could be used to help identify the specific targets of Mac1 during an infection, which will lead to a better understanding of the mechanisms used by mammalian cells to counter virus infection.

## METHODS

### Plasmids

MERS-CoV Mac1 (residues 1110-1273 of pp1a) and mutations were cloned into pET21a+ with a C-terminal His tag. SARS-CoV-2 Mac1 (residues 1023-1197 of pp1a) was cloned into the pET30a+ expression vector with an N-terminal His tag and a TEV cleavage site (Synbio). PARP10-CD was cloned into a pGEX4T expression vector with an N-terminal GST tag and was previously described (kindly provided by Dr. Ivan Ahel, University of Oxford) (52).

### Protein Expression and Purification

A single colony of *E. coli* cells BL21 C41 (DE3) or pRARE (DE3) containing plasmids harboring the constructs of the macrodomain proteins was inoculated into 10 mL LB media and grown overnight at 37°C with shaking at 250 rpm. For most proteins, the overnight culture was transferred to a shaker flask containing TB media at 37°C until the OD600 reached 0.7. The proteins were either induced with either 0.4 mM (SARS-CoV-2 proteins) or 0.05 mM (MERS-CoV proteins) IPTG at 17°C for 20 hours. Cells were pelleted at 3500 × g for 10 min and frozen at -80°C. Frozen cells were thawed at room temperature, resuspended in 50 mM Tris (pH 7.6), 150 mM NaCl, and sonicated using the following cycle parameters: Amplitude: 50%, Pulse length: 30 seconds, Number of pulses: 12, while incubating on ice for >1min between pulses. The soluble fraction was obtained by centrifuging the cell lysate at 45,450 × g for 30 minutes at 4°C. The expressed soluble proteins were purified by affinity chromatography using a 5 ml prepacked HisTrap HP column on an AKTA Pure protein purification system (GE Healthcare). The fractions were further purified by size-exclusion chromatography (SEC) with a Superdex 75 10/300 GL column equilibrated with 20mM Tris (pH 8.0), 150 mM NaCl and the protein sized as a monomer relative to the column calibration standards. For the SARS-CoV-2 N1062A protein several modifications to this protocol were made to obtain stable soluble protein. First, the overnight culture was transferred to LB instead of TB and grown to OD600 0.5 before the protein was induced with 0.05 mM IPTG at 17°C for 20 hours. Cells were resuspended in water prior to sonication. Tris and NaCl were added after sonication. The cell lysate was then incubated with HIS-select HF Nickel Affinity Gel (Millipore-Sigma) overnight, rotating at 4°C. The lysate was then passed into gravity flow chromatography. Columns were washed with 0.5M NaCl and 50 mM Tris-Cl pH 8 and eluted with 0.5 ml of elution buffer with 0.1 M of Imidazole. Following elution, the protein was immediately purified by size-exclusion chromatography as described above. The expression of PARP10 CD protein was previously described (7).

### Isothermal Titration Calorimetry

All ITC titrations were performed on a MicroCal PEAQ-ITC instrument (Malvern Pananalytical Inc., MA). All reactions were performed in 20 mM Tris pH 7.5, 150 mM NaCl using 100 μM of all macrodomain proteins at 25°C. Titration of 2 mM ADP-ribose or ATP (MilliporeSigma) contained in the stirring syringe included a single 0.4 μL injection, followed by 18 consecutive injections of 2 μL. Data analysis of thermograms was analyzed using one set of binding sites model of the MicroCal ITC software to obtain all fitting model parameters for the experiments. MERS-CoV and SARS-CoV-2 WT protein ITC data was previously published (7). These experiments were performed alongside the mutant proteins, thus serving as appropriate controls.

### Differential Scanning Fluorimetry (DSF)

Thermal shift assay with DSF involved use of LightCycler® 480 Instrument (Roche Diagnostics). In total, a 15 μL mixture containing 8X SYPRO Orange (Invitrogen), and 10 μM macrodomain protein in buffer containing 20 mM Hepes, NaOH, pH 7.5 and various concentrations of ADP-ribose were mixed on ice in 384-well PCR plate (Roche). Fluorescent signals were measured from 25 to 95 °C in 0.2 °C/30-s steps (excitation, 470-505 nm; detection, 540-700 nm). Data evaluation and Tm determination involved use of the Roche LightCycler® 480 Protein Melting Analysis software, and data fitting calculations involved the use of single site binding curve analysis on Graphpad Prism.

### AlphaScreen (AS) Assay

The AlphaScreen reactions were carried out in 384-well plates (Alphaplate, PerkinElmer, Waltham, MA) in a total volume of 40 μL in buffer containing 25 mM HEPES (pH 7.4), 100 mM NaCl, 0.5 mM TCEP, 0.1% BSA, and 0.05% CHAPS. All reagents were prepared as 4X stocks and 10 μL volume of each reagent was added to a final volume of 40 μL. All compounds were transferred acoustically using ECHO 555 (Beckman Inc) and preincubated after mixing with purified His-tagged macrodomain protein (250 nM) for 30 min at RT, followed by addition of a 10 amino acid biotinylated and ADP-ribosylated peptide [ARTK(Bio)QTARK(Aoa-RADP)S] (Cambridge peptides) (625 nM). After 1h incubation at RT, streptavidin-coated donor beads (7.5 μg/mL) and nickel chelate acceptor beads (7.5 μg/mL); (PerkinElmer AlphaScreen Histidine Detection Kit) were added under low light conditions, and plates were shaken at 400 rpm for 60 min at RT protected from light. Plates were kept covered and protected from light at all steps and read on BioTek plate reader using an AlphaScreen 680 excitation/570 emission filter set. For data analysis, the percent inhibition was normalized to positive (DMSO + labeled peptide) and negative (DMSO + macrodomain + peptide, no ADPr) controls. The IC_50_ values were calculated via four-parametric non-linear regression analysis constraining bottom (=0), top (=100), & Hillslope (=1) for all curves.

### MAR Hydrolase Assays

First, a 10 μM solution of purified PAPR10-CD protein was incubated for 20 minutes at 37°C with 1 mM final concentration of β-Nicotinamide Adenine Dinucleotide (β NAD^+^) (Millipore-Sigma) in a reaction buffer (50 mM HEPES, 150 mM NaCl, 0.2 mM DTT, and 0.02% NP-40). MARylated PARP10 was aliquoted and stored at -80°C. Next, a 0.5 (I-A/F-A) or 5 (N/A) μM solution of MARylated PARP10-CD and 0.1 (I-A/F-A) or 1 (N-A) μM purified Mac1 protein was added in the reaction buffer (50 mM HEPES, 150 mM NaCl, 0.2 mM DTT, and 0.02% NP-40) and incubated at 37 °C for indicated times. The reaction was stopped with addition of 2X Laemmli sample buffer containing 10% β-mercaptoethanol. Protein samples were heated at 95°C for 5 minutes before loading and separated onto SDS-PAGE cassette (Thermo Fisher Scientific Bolt™ 4-12% Bis-Tris Plus Gels) in MES running buffer. For direct protein detection, the SDS-PAGE gel was stained using InstantBlue® Protein Stain (Expedeon). For immunoblotting, the separated proteins were transferred onto polyvinylidene difluoride (PVDF) membrane using iBlot™ 2 Dry Blotting System (ThermoFisher Scientific). The blot was blocked with 5% skim milk in PBS containing 0.05% Tween-20 and probed with anti-mono ADP-ribose binding reagent MABE1076 (α-MAR) (Millipore-Sigma) and anti-GST tag monoclonal antibody MA4-004 (ThermoFisher Scientific). The primary antibodies were detected with secondary infrared anti-rabbit and anti-mouse antibodies (LI-COR Biosciences). All immunoblots were visualized using Odyssey^®^ CLx Imaging System (LI-COR Biosciences). The images were quantitated using Image J (National Institutes for Health (NIH)) or Image Studio software.

### Molecular Dynamics (MD) Simulations

25 ns simulations were performed for WT and I1153A protein in the presence and absence of ADP-ribose using GROMACS 2019.4 (53). Protein structures used were ADP-ribose-bound SARS-2-CoV Mac1, PDB 6W02 (46), and unbound SARS-2-CoV Mac1, PDB 7KQO (54). The simulations were prepared, including virtual mutagenesis, using CHARMM-GUI’s Solution Builder (55), which was used to build a solvated, rectangular box around one protein, parameterize the ligand, add ions to neutralize the system, set up periodic boundary conditions, and generate the files to perform a gradient based minimization, 100 ps equilibration with a NVT ensemble, and then a 25 ns production run with an NPT ensemble at 303.15 K.

### Cell Culture and Reagents

Vero E6, Huh-7, Vero81, DBT, L929, HeLa cells expressing the MHV receptor carcinoembryonic antigen-related cell adhesion molecule 1 (CEACAM1) (HeLa-MHVR), Baby Hamster Kidney cells expressing the mouse virus receptor CEACAM1 (BHK-MVR) (all gifts from Stanley Perlman, University of Iowa) and A549-ACE2 cells (a gift from Susan Weiss, University of Pennsylvania), were grown in Dulbecco’s modified Eagle medium (DMEM) supplemented with 10% fetal bovine serum (FBS). Calu-3 cells (ATCC) were grown in MEM supplemented with 20% FBS. Bone marrow-derived macrophages (BMDMs) sourced from PARP12^+/+^ and PARP12^-/-^ mice were differentiated into M0 macrophages by incubating cells in Roswell Park Memorial Institute (RPMI) media supplemented with 10% FBS, sodium pyruvate, 100 U/ml penicillin and 100 mg/ml streptomycin, L-glutamine, M-CSF (Genscript) for six days. Then to differentiate into M2 macrophages, IL-4 (Peprotech Inc.) was added for 1 day. Cells were washed and replaced with fresh media every other day after the 4^th^ day. Human IFN-γ was purchased from R&D Systems. Cells were transfected with either Polyjet (Amgen) or Lipofectamine 3,000 (Fisher Scientific) per the instructions of the manufacturers.

### Generation of Recombinant pBAC-JHMV, pBAC-MERS-CoV, and pBAC-SARS-CoV-2 Constructs

All recombinant pBAC constructs were created using Red recombination with several previously described CoV BACs as previously described (41). These include the WT-SARS-CoV-2 BAC based off the Wuhan-Hu-1 isolate provided by Sonia Zuñiga, Li Wang, Isabel Sola and Luis Enjuanes (CNB-CSIC, Madrid, Spain) (56), a MERS-CoV mouse-adapted BAC (a gift from Dr. Stanley Perlman) with GFP inserted into ORF5 (57, 58), and an MHV BAC based off of the JHMV isolate (26). Primers used to create each mutation are listed in Table 2.

**Table 2.**
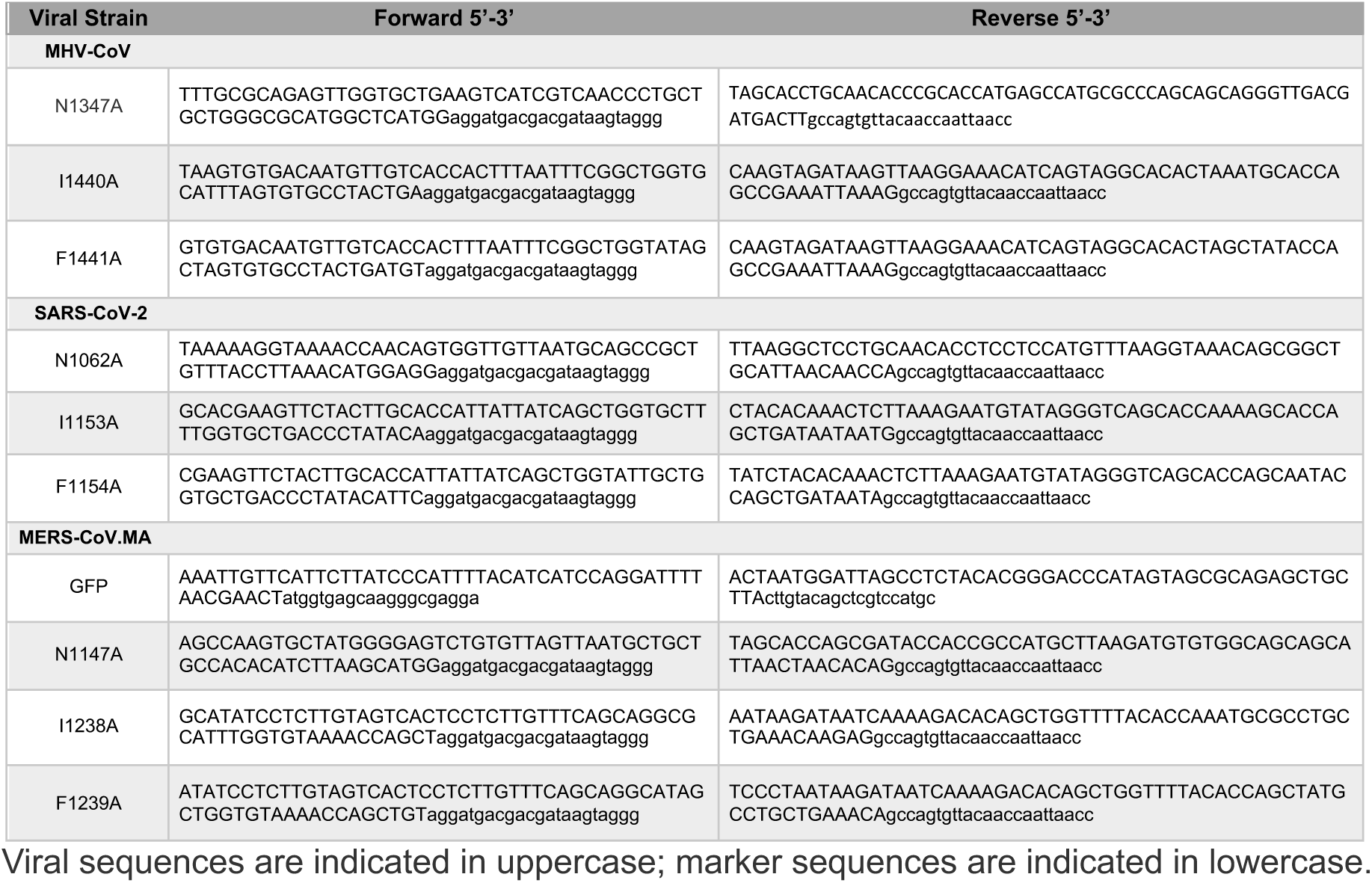
Primers for generating recombinant CoV BACs.

### Reconstitution of Recombinant pBAC-JHMV, pBAC-MERS-CoV, and pBAC-SARS-CoV-2-Derived Virus

All work with SARS-CoV-2 and MERS-CoV was conducted in either the University of Kansas or the Oklahoma State University EHS-approved BSL-3 facilities. To generate SARS-CoV-2 or MERS-CoV, approximately 5 × 10^5^ Huh-7 cells were transfected with 2 μg of purified BAC DNA using Lipofectamine 3,000 (Fisher Scientific) as a transfection reagent. SARS-CoV-2 generated from these transfections (p0) was then passaged in Vero E6 (SARS-CoV-2) or Vero 81 (MERS-CoV) cells to generate viral stocks (p1). All p1 stocks were again sequenced by Sanger sequencing to confirm that they retained the correct mutations. To generate MHV-JHM, approximately 5 × 10^5^ BHK-MVR cells were transfected with 1 μg of purified BAC DNA and 1 μg of N-protein expressing plasmid using PolyJetTM Transfection Reagent (SignaGen). All I-A and F-A recombinant virus stocks had their full genomes sequenced by either Illumina RNA sequencing using polyA enrichment (MHV and MERS-CoV) or by creating amplicons and performing standard whole genome sequencing (SARS-CoV-2) (SeqCenter). No additional variants were detected in these genomes. Sequencing results will be deposited along with all other data in FigShare.

### Mice

Pathogen-free C57BL/6NJ (B6) and K18-ACE2 C57BL/6 mice were originally purchased from Jackson Laboratories and mice were bred and maintained in the animal care facilities at the University of Kansas and Oklahoma State University. Animal studies were approved by the Oklahoma State University and University of Kansas Institutional Animal Care and Use Committees (IACUC) following guidelines set forth in the Guide for the Care and Use of Laboratory Animals.

### Virus Infection

Cells were infected at the indicated MOIs. All infections included a 1 hr adsorption phase. Infected cells and supernatants were collected at indicated time points and titers were determined. For IFN pretreatment experiments, human IFN-γ was added to Calu-3 or A549-ACE2 cells 18 to 20 h prior to infection and was maintained in the culture media throughout the infection. For MHV mouse infections, 5-8 week-old male and female mice were anesthetized with isoflurane and inoculated intranasally with 1×10^4^ PFU recombinant MHV in a total volume of 12 μl DMEM. MHV infected mice were scored for disease based on the following scale: 0: normal, 0-5% weight loss with normal movement and normal behavior; 1: mild disease, 6-12% weight loss, slightly slower movement, and mild neurological issues including circling, sporadic and sudden jumping/hyperreactivity; 2: moderate disease, 13-20% weight loss, slow movement with notable difficulty, moderate neurological issues including occasional circling or head pressing; 3: severe, >20% decrease in weight, severely reduced mobility, and severe neurological symptoms. Mice were euthanized if any of the conditions for a score of 3 were met. For SARS-CoV-2 mouse infections, 12 to 16-wk-old K18-ACE2 C57BL/6 female mice were lightly anesthetized using isoflurane and were intranasally infected with 2.5 × 10^4^ PFU in 50 μL DMEM. To obtain tissue for virus titers, mice were euthanized on different days post challenge, lungs or brains were removed and homogenized in phosphate buffered saline (PBS), and titers were determined by plaque assay on either Hela-MVR (MHV) or Vero E6 (SARS-CoV-2) cells.

### Histopathology

The lung lobes were perfused and placed in 10% formalin. The lung lobes were then processed for H&E staining. The lung lesions were blindly scored by an American College of Veterinary Pathology Board-certified pathologist. The lesions were scored on a scale of 0 to 10% (score 1), 10 to 40% (score 2), 40 to 70% (score 3), and >70% (score 4), and cumulative scores were obtained for each mouse. The lesions scored were bronchointerstitial pneumonia, perivascular inflammation, edema/fibrin, and necrosis.

### Real-time qPCR analysis

RNA was isolated from cells and lungs using TRIzol (Invitrogen) and cDNA was prepared using MMLV-reverse transcriptase as per manufacturer’s instructions (Thermo Fisher Scientific). Quantitative real-time PCR (qRT-PCR) was performed on a QuantStudio3 real-time PCR system using PowerUp SYBR Green Master Mix (Thermo Fisher Scientific). Primers used for qPCR were previously described (24). Cycle threshold (C_T_) values were normalized to the housekeeping gene hypoxanthine phosphoribosyltransferase (HPRT) by the following equation: C_T_ = C_T(gene of interest)_ - C_T(HPRT)_. Results are shown as a ratio to HPRT calculated as 2^-ΔCT^.

### Statistics

A Student’s *t* test was used to analyze differences in mean values between 2 groups, for multiple group comparisons, a one-way ANOVA was used. All results are expressed as means ± standard errors of the means (SEM) unless stated as standard differentiation (SD). P values of ≤0.05 were considered statistically significant (*, P≤0.05; **, P≤0.01; ***, P≤0.001; ****, P ≤0.0001; ns, not significant).

## ACKNOWLEDGEMENTS

We thank Ivan Ahel for providing protein expression plasmids, Stanley Perlman and Susan Weiss for cell lines, Stanley Perlman for critical reading of the manuscript and the MERS-CoV mouse adapted BAC, and Luis Enjuanes and Sonia Zuñiga for the SARS-CoV-2 BAC. We thank Brian Sanderson, the KU Center for Genomics, and the K-INBRE Genomic Data Science Core supposted by the IDeA Program of the NIGMS award number P20GM103418 for help with RNAseq analysis. We thank our funding from the NIH, an NIH Graduate Training grant, the University of Kansas College of Liberal Arts and Sciences, and the University of Kansas Madison and Lila Self graduate programs.

## Funding

National Institutes of Health (NIH) grant R35GM138029 (ARF)

National Institutes of Health (NIH) grant P20GM113117 (ARF)

National Institutes of Health (NIH) grant K22AI134993 (ARF)

National Institutes of Health (NIH) grant P20GM103648 (RC)

NIH Graduate Training at the Biology-Chemistry Interface grant T32GM132061 (CMK) University of Kansas College of Liberal Arts and Sciences Graduate Research Fellowship (CMK)

University of Kansas Madison and Lila Self Scholarship and Fellowship (JJP)

The funders had no role in study design, data collection and analysis, decision to publish, or preparation of the manuscript.

## Author contributions

Conceptualization: CMK, JJP, YMA, ARF

Data curation: CMK, YMA, AR, JJOC, RG, SM, RC, ARF

Formal analysis: CMK, YMA, AR, JJOC, RG, SM, RC, ARF

Funding acquisition: CMK, RC, ARF

Methodology: CMK, JJP, YMA, AR, JJOC, RG, RK, PG, DKJ, SM, RC, ARF

Investigation: CMK, JJP, YMA, AR, JJOC, RG, RS, PS, SP, PM, RK, DKJ, SM, RC, ARF

Project administration: RC, ARF

Resources: AR, PG, RC, ARF

Visualization: CMK, JJP, YMA, AR, JJOC, RG, RS, PM, RK, DKJ, SM, RC, ARF

Validation: CMK, AR, SM, RC, ARF

Supervision: PG, RC, ARF

Writing—original draft: CMK, ARF

Writing—review & editing: CMK, JJP, YMA, AR, JJOC, RG, RS, PS, SP, PM, RK, DKJ, SM, RC, ARF

The authors have no competing interests.

## Notes

### Competing Interest Statement

The authors have declared no competing interest.

### Summary of Updates

Figures 5 and 6 were revised, with figure 6 now being the combination of two prior figures. Several changes to the text and title of the manuscript were also made. New authors were added.

